# L-pentahomoserine correlates with therapy outcome in esophageal cancer and promotes metabolic adaptations that support cell survival under nutrient-deprived conditions

**DOI:** 10.64898/2026.01.15.699640

**Authors:** Mark E. Kelly, Alejandro Huerta-Uribe, Michaela Cerná, Keira B. Hughes, Hasnain Ahmed, Marco F. Fernandes, Kevin M. Rattigan, Ruhi Deshmukh, David S. Moura, Swati Arya, Stephanie May, Miryam Müller, Thomas M. Drake, Anastasia Georgakopoulou, Thomas G. Bird, Michael Hisbergues, Guillaume Piessen, Alan J. Stewart, Allan J. B. Watson, David Sumpton, Victor H. Villar

**Affiliations:** School of Medicine, University of St Andrews, North Haugh, KY16 9TF, UK; Cancer Research UK Scotland Institute; Garscube Estate, Switchback Road, Glasgow, G61 1BD, UK; EaStCHEM, School of Chemistry, University of St Andrews, North Haugh, St Andrews KY16 9ST, UK; Wolfson Wohl Cancer Research Centre; Institute of Cancer Sciences, University of Glasgow, Glasgow G61 1QH, UK; Instituto de investigación Sanitaria Fundación Jiménez Diaz (IIS/FJD; UAM), Madrid, Spain; Centre for Inflammation Research, University of Edinburgh, Edinburgh, UK; University Lille, Inserm, CHU Lille, Biological Resource Center (CRB-CIC1403), F-59037 Lille CEDEX, France; University Lille, CNRS, Inserm, CHU Lille, UMR9020-U1277 - CANTHER - Cancer Heterogeneity, Plasticity and Resistance to Therapies, F-59000, Lille, France

## Abstract

Esophageal adenocarcinoma (EAC) is the sixth-leading cause of cancer-related death. Although pyrimidine analogue-based neoadjuvant and adjuvant therapies are widely used, patient responses remain variable. Emerging evidence indicates that bacteria-derived metabolites influence tumor biology and therapy outcomes. To identify non-canonical plasma metabolites linked to cancer biology, we performed correlation analyses between untargeted metabolomics profiles and overall survival. This approach revealed a bacterial metabolite called L-pentahomoserine, or L-2-amino-5-hydroxypentanoic acid (L-2A5HPA), to be positively associated with overall survival. Notably, L-2A5HPA promoted cell survival under nutrient limitation by redirecting glucose metabolism towards aspartate and pyrimidine biosynthesis. In vitro, L-2A5HPA uptake varied among cell lines and was controlled by stereospecific transporters. Furthermore, metabolic profiling in mouse models of liver cancer showed different levels of L-2A5HPA and a strong correlation with pyrimidine intermediates, dihydroorotate and orotate. The link between L-2A5HPA, pyrimidine nucleotide metabolism, and cell survival provides mechanistic insight into its association with patient outcome. Our findings position L-2A5HPA as a metabolite with potential to become a prognostic biomarker for EAC and underscores its role in metabolic adaptation under nutrient-deprived conditions.

## Introduction

Esophageal cancer (EC) is the seventh most common cancer worldwide and the sixth-leading cause of cancer related deaths^1,2^. The most common subtypes of ECs are esophageal squamous cell carcinoma (ESCC) and esophageal adenocarcinoma (EAC). Although these subtypes are epidemiologically and biologically very distinct, both share a poor outcome with a five-year survival rate of 10%^3,4^. ESCC accounts for 90% of EC cases, and is more prevalent in East Africa and South America, whereas EAC is more frequent in Europe, America, and Australia^3,5,6^. Whilst ESCC remains the dominant form globally, EAC incidence rates are increasing sharply in developed countries^3^. EAC typically arises from Barrett’s esophagus, a preneoplastic condition characterized by columnar intestinal-type mucosa, which is the main risk factor for the development of EAC^7,8^.

The therapeutic strategy for EC depends on the tumor stage. Early-stage disease can be treated with endoscopic resection, whereas locally advanced stages of EC are treated with neoadjuvant chemotherapy or radio-chemotherapy, followed by surgery or exclusive radio-chemotherapy^9^. Although there is evidence showing that preoperative chemotherapy or radio-chemotherapy provides a survival benefit for patients compared with surgery alone^10^, a variable treatment response has been observed^10–14^. Diet and gut bacterial-derived metabolites have attracted significant attention due to their impact on health and disease^15^. These metabolites have shown to influence tumor formation and progression, as well as responses to chemotherapy, radiotherapy, and immunotherapy, explaining potentially the variable responses^16–19^. The identification of non-canonical circulating metabolites could uncover bacteria-derived metabolites capable of predicting chemotherapy outcome and modulate tumor biology, enhancing patient stratification and therapy selection.

Herein we report that a bacterial amino acid called L-pentahomoserine or L-2-amino-5-hydroxypentanoic acid (L-2A5HPA), showed association with therapy response and is enriched in the plasma of EAC patients who respond favorably to chemotherapy. Furthermore, we demonstrate that L-2A5HPA supports cancer cell survival under nutrient limitation by redirecting glucose metabolism towards aspartate and pyrimidine synthesis. Our findings highlight the potential of L-2A5HPA as a prognostic metabolite in esophageal adenocarcinoma and uncovers its novel role in tumor biology.

## Results

### C_5_H_11_NO_3_ plasma levels are associated with therapeutic outcome in esophageal adenocarcinoma patients

To identify metabolites that can be associated with chemotherapy efficacy and cancer biology, we performed a correlation analysis between the overall survival data available from eleven esophageal adenocarcinoma patients (FREGAT clinical-biological database) and their pre-treatment plasma level of all the metabolites detected by untargeted metabolomic analysis. These patients, who were all males were treated with pyrimidine analogues-containing therapy, including 5-fluorouracil-based chemotherapy regimens such as Folfox (oxaliplatin, leucovorin, 5-fluorouracil, 81.8%) and ECX (epirubicin, cisplatin, capecitabine,18.2%). The patient characteristics are presented in **Extended Data Figure 1a**.

All metabolites detected through untargeted analysis were plotted based on their Pearson correlation coefficient and average peak area (a key parameter for structure elucidation), against the corresponding –log *P* values derived from the Pearson correlation analysis (**Figure 1a**). After filtering those with retention time (RT) higher than 2 mins to avoid poor retained metabolites and ion suppression effects, we identified metabolites that exhibited a positive correlation with the overall survival of the patients. The five metabolites that showed the strongest correlations were C_8_H_11_NO_6_S (m/z=250.0377 RT=11.812 min., positive mode), C_5_H_7_NO_2_ (m/z=114.0549, RT=2.619 min., positive mode), C_6_H_10_O_3_ (m/z=148.0967, RT=6.964 min., positive mode), C_5_H_11_NO_3_ (m/z=134.0809, RT=8.055 min., positive mode), C_6_H_6_O_5_S (m/z=188.9862, RT=2.174 min., negative mode) (**Figure 1a**). The most abundant metabolite that displayed both a high Pearson coefficient and an appropriate retention time for our analytical method was C_5_H_11_NO_3_. This metabolite showed a strong correlation with patient survival (Pearson r=0.8495, *P*=0.0009) (**Figure 1b**). In addition, the stratification of patients by C_5_H_11_NO_3_ levels into higher or equal, and lower than the median, showed also a significant difference in survival between the groups (**Figure 1c**, *P*=0.0007). Therefore, assessing pre-treatment plasma levels of C_5_H_11_NO_3_ may offer a valuable tool for predicting overall survival in EAC patients.

**Figure 1:**
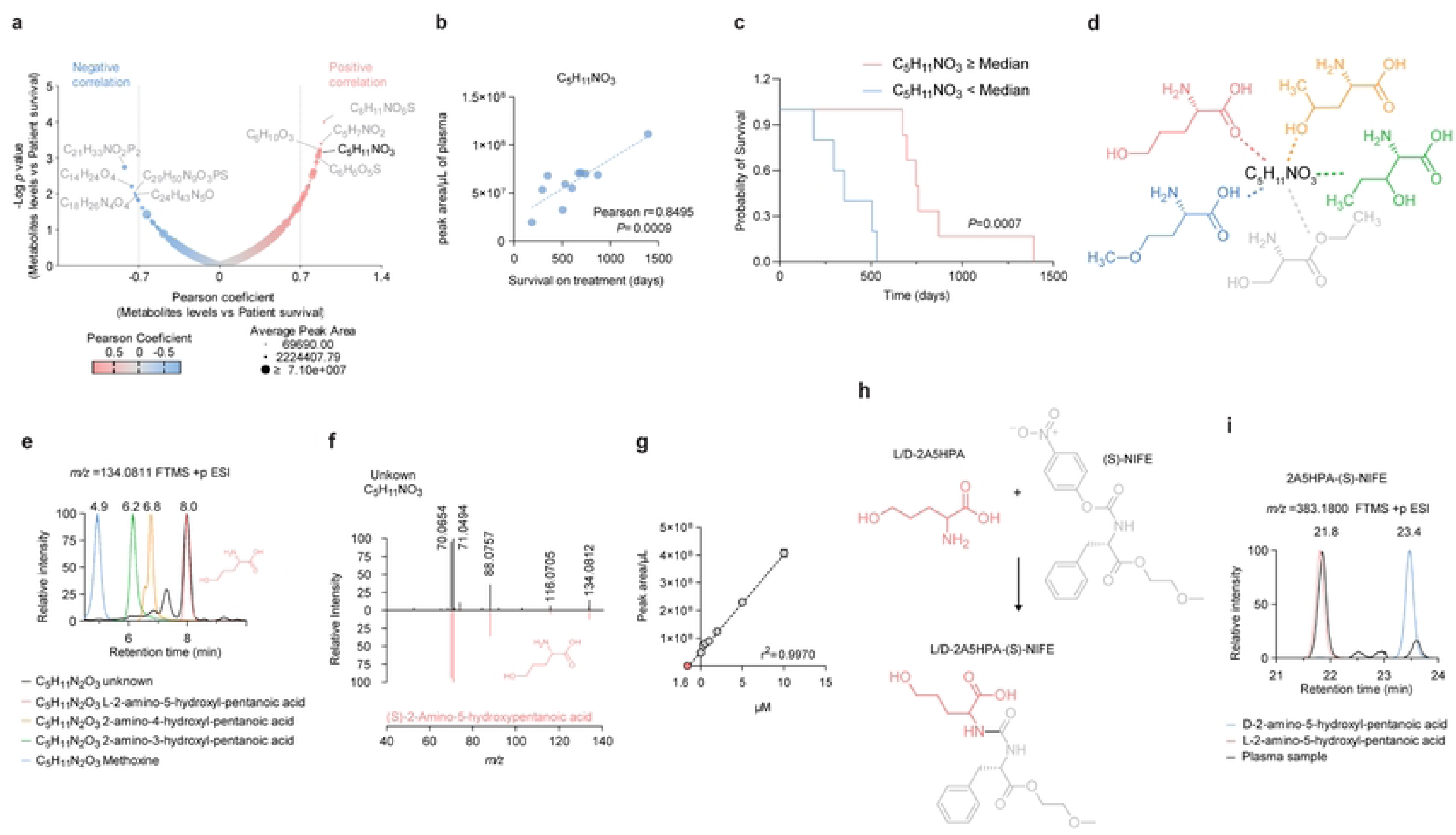
Identification and structure elucidation of C_5_H_11_NO_3_ as L-2-amino-5-hydroxypentanoic acid, a novel metabolite associated with therapeutic outcome in esophageal adenocarcinoma. **a**) Bubble plot showing the correlation between all the detected metabolites and the overall survival of the patients (n=11). Bubble size reflects the abundance of the metabolites (average peak area). Positive and negative correlations are shown in red and blue, respectively. Metabolites highlighted in grey were discarded as they either have low abundance or their retention time is not appropriate for our method. C_5_H_11_NO_3_ is highlighted in black as the selected metabolite to elucidate. **b**) Correlation between the plasma levels of C_5_H_11_NO_3_ and the overall survival of the eleven esophageal cancer patients. Pearson r values and the *P* values are shown in the graph. **c**) Kaplan–Meier curve showing survival differences between patients stratified by plasma levels of C5H11NO3. Patients with levels greater than or equal to the median (red), and those with lower levels (blue) are presented. The p-value from a Logrank (Mantel-Cox) test is shown on the graph. **d**) Different potential chemical structures for C_5_H_11_NO_3_. **e**) Extracted ion chromatogram (XIC) for the unknown metabolite from pooled patient plasma (black), L-2-amino-5-hydroxy-pentanoic acid (L-2A5HPA, pentahomoserine, red), 2-amino-4-hydroxyl-pentanoic acid (orange), 2-amino-3-hydroxyl-pentanoic acid (green), methoxine (blue). **f**) Mirror plot showing the fragmentation pattern of the unknown metabolite from pooled plasma samples and chemically pure standard of L-2A5HPA. **g**) Absolute quantification 2A5HPA using the standard addition method in pooled patient plasma (n=4). The intercept on X is highlighted in red, which represents the plasma concentration of L-2A5HPA. **h)** Schematic showing the reaction between both enantiomers L-2A5HPA and D-2A5HPA and (S)-NIFE. **i**) Extracted ion chromatogram (XIC) showing the (S)-NIFE-derivatized L-2A5HPA standard (red), (S)-NIFE-derivatized D-2A5HPA standard (blue) and (S)-NIFE-derivatized 2A5HPA from pooled plasma (black). Circles represent individual metabolites, patients or the average of technical duplicates.

### C_5_H_11_NO_3_ identification as L-pentahomoserine

Given the potential for C_5_H_11_NO_3_ levels to predict overall survival, we focused on the elucidation of its chemical structure. Several molecules have this chemical formula. Hence, we systematically tested several possible compounds, including methoxine, ethylserinate, 2-amino-3-hydroxypentanoic acid, 2-amino-4-hydroxypentanoic acid, and 2-amino-5-hydroxypentanoic acid (**Figure 1d**). Based on the retention time of the unknown metabolite (RT=8.055 min.), we were able to discard many of the potential alternative structures for C_5_H_11_NO_3_. Methoxine exhibited a RT=4.9 min, ethyl-serinate RT=1.7 and 2.5 min., 2-amino-3-hydroxypentanoic acid RT=6.2 min., and 2-amino-4-hydroxypentanoic acid RT=6.8 min. (**Figure 1e**). However, 2-amino-5-hydroxypentanoic acid (2A5HPA) or L-pentahomoserine showed the same RT and fragmentation pattern as the unknown molecule, exhibiting common fragments with a *m/z* of 70.0654, 71.0494, 88.0757, and 116.0705, which were used to identify the metabolite as 2A5HPA (**Figure 1e and 1f**). In addition, to further corroborate the identity of the unknown metabolite, we compared the fragmentation pattern of the unknown metabolite with ethyl-serinate (**Extended Data Figure 1b**), 2-amino-3-hydroxypentanoic acid (**Extended Data Figure 1c),** 2-amino-4-hydroxypentanoic acid (**Extended Data Figure 1d),** and methoxine (**Extended Data Figure 1e**). All of which exhibited different fragmentation patterns, further refuting these as possible candidates for C_5_H_11_NO_3_.

### Estimation of the physiological concentrations of 2A5HPA in plasma and tissue, and identification of the predominant enantiomer in human plasma

To obtain information about the physiological concentration of 2A5HPA, we performed a standard addition method to quantify the absolute concentration of 2A5HPA in human plasma. The concentration of 2A5HPA in a pooled plasma sample from patients was ∼1.6 µM (n=4, **Figure 1g**). Furthermore, we also measured the levels of 2A5HPA in pooled mouse plasma (n=3), which was ∼1.1 µM (**Extended Data Figure 1f**). To obtain an approximation of the physiological concentration of 2A5HPA in tissue, we examined mouse esophagus tissue (n=1) where the concentration was ∼167 µM (**Extended Data Figure 1g**). Similar levels were found in mouse liver (∼164 µM, n=1) (**Extended Data Figure 1h**). The difference in the concentration of 2A5HPA between mouse plasma and mouse tissue suggests a potential active mechanism for its uptake or an increased retention in the tissues.

Different enantiomeric forms of amino acids can lead to different biological effects ^20^, we decided to define the stereochemistry of 2A5HPA. We chemically synthesized both the D and L enantiomers of 2A5HPA ^21–23^ (**Extended Data Figure 1i**). To distinguish both enantiomers qualitatively, we employed a modified (S)-NIFE method, which enables the separation of enantiomers without a chiral chromatographic column^24,25^ (**Figure 1h**). The different retention times observed for the (S)-NIFE-derivatized L-2A5HPA (RT=21.8 min.) and (S)-NIFE-derivatized D-2A5HPA (RT=23.4 min.) allowed us to conclude that the L-2A5HPA enantiomer is the predominant form in human plasma samples (**Figure 1i**). We further confirmed the formation of these products by identifying the fragments of D- and L-2A5HPA in a LC-MS/MS analysis (**Extended Data Figure 1j and 1k**).

### L-2A5HPA promotes cell survival under nutrient-deprived conditions

The strong correlation between plasma levels of L-2A5HPA and overall patient survival suggests a potential role for this metabolite in cancer biology. Given its structural similarity with serine, we hypothesized that L-2A5HPA could alter serine metabolism (**Extended Data Figure 2a**). To test this hypothesis, we used two esophageal cancer cell lines, SK-GT-4 and OA-P4C, and we cultured them with physiological media (Plasmax^TM^) in the presence or absence of L-2A5HPA (50 µM) for 24h. Untargeted metabolomic analysis did not show any intermediates of serine metabolism, or the central carbon metabolism to be altered upon addition of L-2A5HPA in either SK-GT-4 or OA-P4C cell lines, suggesting that this metabolites is not incorporated into the mammalian metabolism (**Extended Data Figure 2b and 2c**).

**Figure 2.**
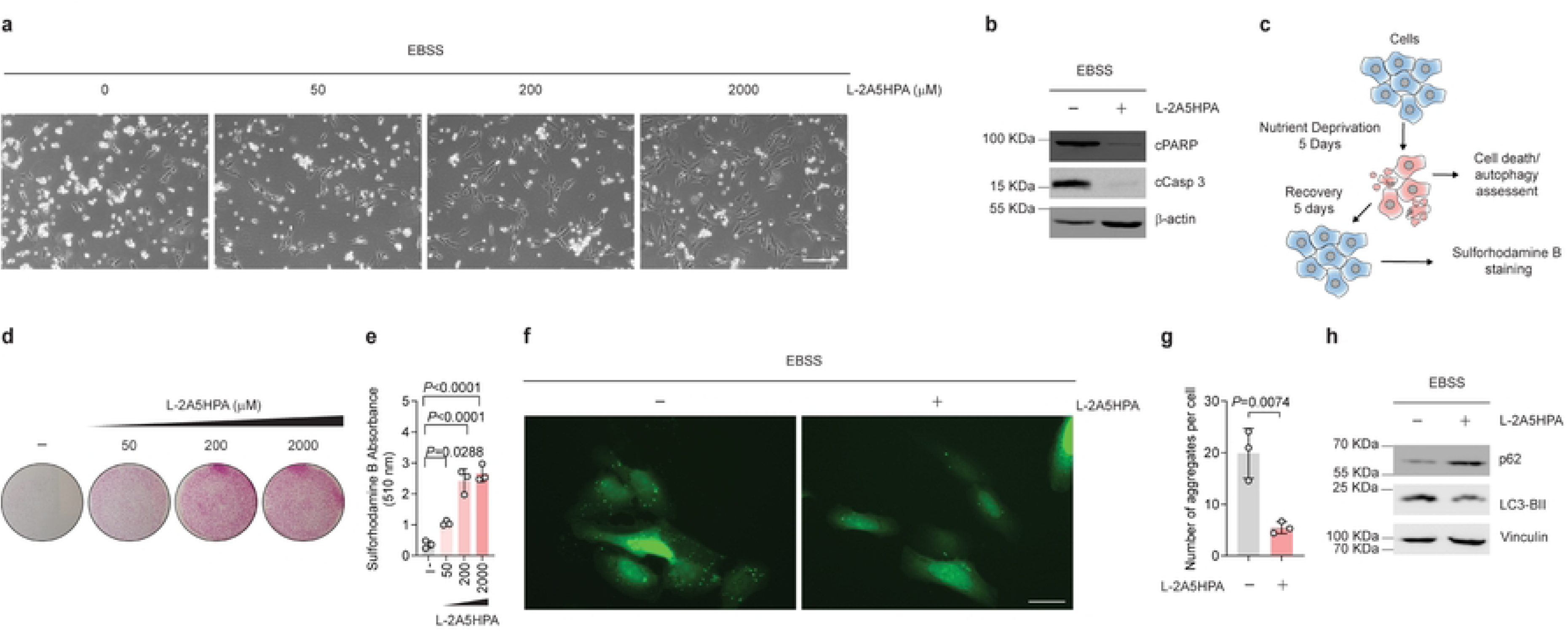
L-2A5HPA promotes survival in conditions of nutrient deprivation. **a**) Representative images of U2OS cells exposed to nutrient deprivation (EBSS) in the presence or absence of L-2A5HPA (0, 50µM, 200 µM and 2mM) for 5 days. Scale bar represents 100 µm. **b**) Immunoblot analysis of apoptosis markers, PARP and caspase 3 in U2OS starved for 5 days in the presence or absence of L-2A5HPA. β-actin was used as a loading control. **c**) Schematic showing the protocol used for starvation and recovery experiments. U2OS cells were starved for 5 days on EBSS, and apoptosis or autophagy was assessed, or cells were allowed to grow in complete media for 5 extra days and stained with sulforhodamine B. **d**) Representative image from cells treated as in **a**), which were then exposed to complete media for 5 days and stained with sulforhodamine B. **e**) Quantification of sulforhodamine B absorbance at 510 nm from three biological replicates. **f**) Representative images from U2OS GFP-LC3 cells in the presence or absence of L-2A5HPA (2 mM) for 5 days. The autophagosome formation of GFP-LC3 aggregates was assessed by fluorescent microscopy ^33^. The scale bar represents 25 µm. **g**) Quantification of the number of aggregates per cell in the conditions described in **f**). **h**) Immunoblot analysis of autophagy markers, p62 and LC3BII in U2OS cells after 5 days on nutrient deprivation with or without 2A5HPA (2 mM). Vinculin was used as a loading control. Bars represent the average ± SEM of three independent replicates. Data was analyzed using a one-way ANOVA or Student’s t-test. Circles represent independent experiments.

In addition, the presence of L-2A5HPA did not alter the intracellular levels of serine in these cell lines (**Extended Data Figure 2d and 2e**), and no correlation between the levels of L-2A5HPA and serine was observed in the plasma samples from patients (Pearson r=-0.2530 *P*=0.4528, **Extended Data Figure 2f**). This evidence supports the notion that L-2A5HPA does not alter serine metabolism or compete for its uptake by transporters.

The high proliferation of tumour cells often leads to nutrient deprived conditions ^26,27^, which can be exacerbated in esophageal cancer patients who are prone to developing malnutrition due to the location of the tumors or chemotherapy ^28,29^. In addition, the accumulation of L-2A5HPA in bacteria appears to be also linked with nutrient deprivation conditions^3,30–32^. Therefore, the potential biological effect of L-2A5HPA in nutrient-deprived conditions was explored. We investigated the effect of L-2A5HPA on the survival of cancer cells and autophagy using the parental U2OS cells and U2OS GFP-LC3, which constitutively express an autophagy reporter^33,34^. Unexpectedly, while U2OS cells exposed to nutrient deprivation conditions (EBSS) underwent cell death after 5 days of treatment, cells treated with L-2A5HPA remained viable in a concentration-dependent manner (0, 50 µM, 200 µM and 2 mM, **Figure 2a**). To further test whether L-2A5HPA prevents apoptotic cell death, we measured the apoptosis markers cleaved PARP and cleaved caspase 3 by western blotting. L-2A5HPA supplementation prevented the activation/cleavage of both caspase 3 and PARP that is induced in nutrient deprivation conditions (**Figure 2b**). Furthermore, when cells were starved and then switched to complete media for 5 days, recovery conditions (**Figure 2c**), we observed that cells treated with L-2A5HPA recovered and grew faster than the vehicle-treated cells (**Figure 2d and 2e**). The ability of L-2A5HPA to promote survival of cells upon nutrient deprivation was also observed in another cell line (synovial sarcoma SW982, **Extended Data Figure 2g and 2h**). It is known that certain amino acids, such as glutamine and leucine, can induce cell death in these conditions by activating mTORC1 and inhibiting autophagy^33,34^. Therefore, we assessed whether L-2A5HPA can modulate the mTORC1/autophagy pathway in conditions of nutrient deprivation. Indeed, L-2A5HPA activated mTORC1 signaling assessed by phospho-S6 (S235/236) (**Extended Data Figure 2i**). As a consequence, L-2A5HPA also inhibited autophagy, as determined by the decrease in the number of GFP-LC3 aggregates per cell (U2OS GFP-LC3 ^33,34^, the increase in p62 and the decrease in LC3BII levels (**Figure 2f-h**).

Despite L-2A5HPA and L-serine sharing similar chemical structures, L-serine significantly promoted the recovery of the cells after starvation, although its effect was less pronounced than that of L-2A5HPA (**Extended Data Figure 2j-l**). Furthermore, we also examined whether the pro-survival effect of 2A5HPA was stereospecific. Interestingly, only the L-enantiomer of 2A5HPA was able to protect the cells under nutrient-deprived conditions, which suggests a highly specific biological role for this metabolite (**Extended Data Figure 2m-o**). To gain insight into this specificity, we assessed the uptake of both enantiomers. The uptake of L-2A5HPA was significantly more efficient than that of D-2A5HPA (**Extended Data Figure 2p**), which can explain the inability of D-2A5HPA to support cell survival (**Figure 2m-o**). These findings suggest that the stereospecific effect of 2A5HPA is likely mediated by transporters that can distinguish between enantiomers.

### L-2A5HPA induces a diversion of glucose metabolism towards the synthesis of aspartate and pyrimidines, promoting survival upon nutrient-deprived conditions

Although L-2A5HPA supplementation inhibited autophagy, which is a survival mechanism activated upon nutrient deprivation ^33,34^, it paradoxically promoted cell survival. Under our nutrient-limiting conditions, glucose serves as the primary nutrient source, suggesting that L-2A5HPA may induce a rewiring of glucose metabolism to sustain survival. To test this hypothesis, we performed ^13^C_6_-glucose tracing for 72h under nutrient-deprived conditions (**Figure 3a**). We observed that the presence of L-2A5HPA, induced a decrease in the enrichment of ^13^C_3_-pyruvate (**Figure 3b**), without affecting ^13^C_3_-lactate (**Extended Data Figure 3a**). Given that the decrease of pyruvate can be explained by both an increase in utilization of pyruvate by the TCA cycle or a diversion to other pathways, we first analyzed the enrichment of TCA cycle metabolites. Whilst the presence of L-2A5HPA decreased the enrichment of ^13^C_2_-αKG (**Figure 3c**) and ^13^C_2_-malate (**Figure 3d**), it also increased the enrichment of ^13^C_2_-succinate (**Figure 3e**) and ^13^C_2_-aspartate (**Figure 3f**). This pattern suggests a decrease in the incorporation of carbons from pyruvate to acetyl-CoA and the TCA cycle, and a potential increase in the synthesis of aspartate from pyruvate. Pyruvate carboxylase (PC) is a mitochondrial enzyme that utilizes pyruvate to produce oxaloacetate, which in turn, is transaminated by glutamate–oxaloacetate transaminase (GOT) into aspartate (**Figure 3a**). Indeed, the enrichment of isotopologues used to assess the activity of PC, ^13^C_3_-aspartate (**Figure 3g**) and ^13^C_5_-αKG (**Figure 3h**)^35^, was significantly increased by the presence of L-2A5HPA. This suggested that aspartate limitation may be critical for survival, as it has been previously reported to be important for the synthesis of pyrimidines and for proliferation^36,37^. Based on this, we also assessed the enrichment of the pyrimidine pathway ^13^C_5_-UMP (from ^13^C_6_-glucose), ^13^C_7_-UMP (from ^13^C_6_-glucose and ^13^C_2_-aspartate) and ^13^C_8_-UMP (from ^13^C_6_-glucose and ^13^C_3_-aspartate). While the enrichment of ^13^C_5_-UMP was decreased (**Figure 3i**), the enrichment of ^13^C_7_-UMP (**Figure 3j**) and ^13^C_8_-UMP (**Figure 3k**) were increased, with a higher contribution from the ^13^C_8_-UMP, which is synthesized from ^13^C_3_-aspartate. This finding provides evidence for the diversion of glucose towards the synthesis of pyrimidines, which is supported by both the synthesis of ribose and aspartate^37^. To assess whether this observation is dependent simply on changes in the metabolic fluxes or is related to the levels of PC, we measured protein levels of PC in these conditions. While cells treated with vehicle or D-2A5HPA exhibited low levels of PC, cells treated with L-2A5HPA had similar levels of PC as non-starved cells (**Figure 3l**). Hence, the presence of L-2A5HPA enables a glucose metabolism diversion towards pyrimidine synthesis, thereby supporting cell survival through maintaining PC expression. To determine whether PC/GOT activity is sufficient to rescue viability in these conditions, we supplemented starved cells with aspartate (2 mM) or pyruvate (2 mM), metabolites that are downstream and upstream of PC/GOT activities, respectively. As predicted, pyruvate was not sufficient to rescue the cell viability, potentially due to the low levels of PC observed on starvation (**Figure 3l**). In contrast, aspartate, the product of PC/GPT activity, was able to rescue the viability and the proliferation of cells (**Figure 3m-o**). This highlights the availability of aspartate and pyrimidine nucleotides to be critical factors for survival in these conditions.

**Figure 3.**
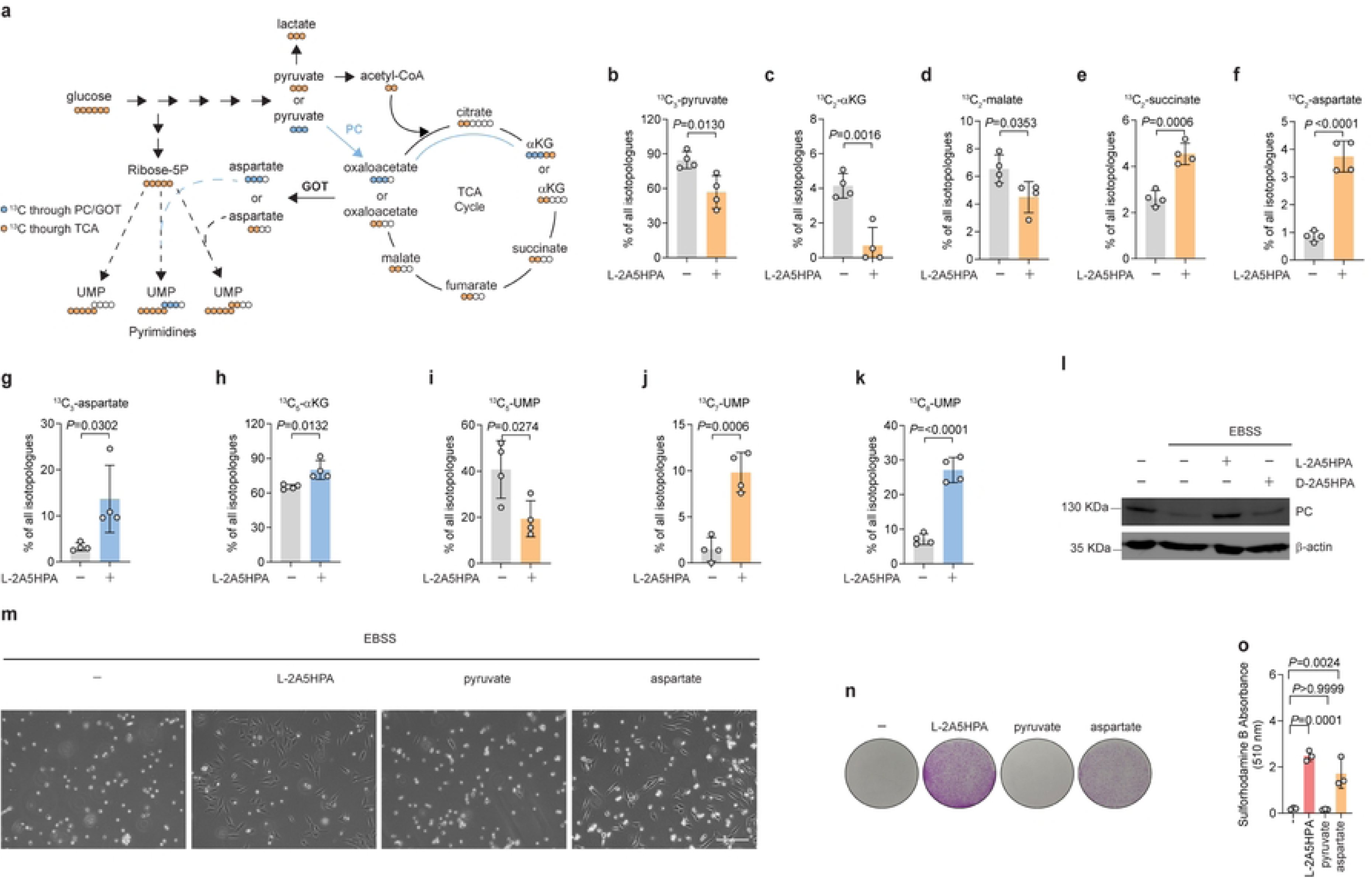
L-2A5HPA diverts glucose metabolism towards the synthesis of aspartate and pyrimidines upon nutrient deprivation. **a**) Schematic depicting glucose metabolism and the distribution of ^13^C-carbons across the different pathways: glycolysis, TCA cycle, amino acids and pyrimidines. **b-k**) Relative levels of ^13^C_2_ or ^13^C_3_ isotopologues from ^13^C_6-_Glucose in U2OS cells upon nutrient deprivation in the presence or absence of L-2A5HPA during 72h. The enrichment of ^13^C_3_-pyruvate (**b**) ^13^C_2_-αKG (**c**), ^13^C_2_-malate (**d**), ^13^C_2_-succinate (**e**), ^13^C_2_-aspartate (**f**), ^13^C_3_-aspartate (**g**), ^13^C_5_-αKG (**h**), ^13^C_5_-UMP (**i**), ^13^C_7_-UMP (**j**), ^13^C_8_-UMP (**k**) are shown. **l**) Immunoblot analysis for pyruvate carboxylase (PC) in U2OS cells in complete media, or upon starvation for 5 days in the presence of L-2A5HPA (2 mM) or D-2A5HPA (2mM). β-actin was used as a loading control. **m**) Representative image of U2OS cells exposed to nutrient deprivation and treated with either vehicle, L-2A5HPA (2 mM), pyruvate (2 mM) or aspartate (2 mM) for 5 days. Scale bar equals 100 µm. **n**) Representative image from cells treated as in **m**), which were then exposed to complete media for 5 days and stained with sulforhodamine B. **o**) Quantification of sulforhodamine B absorbance at 510 nm of three biological replicates. Data were analyzed using a Student’s t-test or one-way ANOVA. Bars represent the average ± SEM of three or four independent replicates. Circles represent independent experiments.

### L-2A5HPA promotes cell survival though an autophagy-independent mechanism

Nutrient deprivation can lead to mitochondrial damage and mitophagy induction ^38^, resulting likely in decreased levels of PC and survival. Given that L-2A5HPA inhibits autophagy, it may also prevent mitochondrial degradation, thereby maintaining higher PC levels, supporting the synthesis of aspartate and pyrimidines, ultimately promoting cell survival. To examine the role of autophagy in the survival phenotypes induced by L-2A5HPA, we reactivated autophagy using rapamycin. Autophagy reactivation did not alter the ability of L-2A5HPA to promote cell survival (**Extended Data Figure 3b-d**). Furthermore, the rescuing effect of aspartate was not linked to autophagy inhibition as the levels of p62 remained similar between aspartate-supplemented and the vehicle-treated conditions (**Extended Data Figure 3e**). Altogether, these findings indicate that the metabolic role of L-2A5HPA is the primary contributor to cell survival under nutrient deprived conditions.

### L-2A5HPA uptake and intratumoural levels are variable

Considering the unexpected in vitro findings alongside the strong positive correlation between L-2A5HPA levels and overall patient survival, we aimed to explore and better understand the underlying mechanism behind these observations. L-2A5HPA is a known bacterial metabolite found in *E. coli* and the microbiome of bees^30,39^. To date, no mammalian pathway that would enable its synthesis or degradation has been described. Thus, we hypothesized that tumor cells uptake L-2A5HPA from plasma and that its biological effect may depend on their ability to uptake this metabolite. To examine this hypothesis, we incubated three cell lines, U2OS, SK-GT-4 and OAC-P4C, with Plasmax^TM^ media containing 50 µM of L-2A5HPA for one minute. We observed that the intracellular levels of L-2A5HPA differed in one of the three cell lines tested (**Extended Data Figure 3f**). In addition, we assessed the levels of L-2A5HPA within the tumor microenvironment of four different genetically engineered mouse models of hepatocellular carcinoma (HCC) ^40^. We observed that two mouse models exhibited significantly higher levels of L-2A5HPA compared with healthy tissue (**Extended Data Figure 3g**). Of note, a correlation analysis of L-2A5HPA and all the metabolites detected by a targeted analysis in healthy liver and tumor tissues (**Extended Data Figure 3h**), identified a strong association with the levels of dihydroorotate (Pearson r= 0.8462 and *P*<0.0001, **Extended Data Figure 3i**) and orotate (Pearson r= 0.7470 and *P*<0.0001, **Extended Data Figure 3j)**, both intermediates of the *de novo* synthesis of pyrimidines. Together, this evidence suggests that the uptake mechanism for L-2A5HPA is not simple diffusion, but an active process likely mediated by transporters, and that the levels of L-2A5HPA are highly associated with the *de novo* synthesis of pyrimidine nucleotide intermediates *in vivo*.

## Discussion

In this study, we identified L-pentahomoserine or L-2-amino-5-hydroxypentanoic acid (L-2A5HPA) as a metabolite showing a positive correlation with EAC patient survival. In addition, L-2A5HPA plasma levels enable stratification with promising ability to predict responses to 5-fluorouracil-containing chemotherapy. Despite the origin of L-2A5HPA being linked to bacterial metabolism, no common biosynthetic pathway has yet been described in mammals^30,39,41^. Accordingly, we did not detect any incorporation of L-2A5HPA into polar metabolism, including central carbon, amino acid or nucleotide metabolism, upon nutrient sufficient conditions. However, there is a report suggesting the incorporation of L-pentahomoserine (L-2A5HPA) into a novel class of phospholipids (PX, named by the authors) in the enteric pathogen *Campylobacter jejuni* upon hypoxic conditions^31^. However, it is not known whether L-2A5HPA can be incorporated or not into lipids in mammals.

The accumulation of L-2A5HPA in bacteria is linked with nutrient deprivation conditions, and given the localisation of the tumour in EAC, patients are more prone to suffer from malnutrition due to decreased food ingestion^3,28,32^. This could highlight a potential role for L-2A5HPA in conditions of nutrient scarcity. Indeed, despite activating mTORC1 signalling and inhibiting autophagy, a process that is crucial for cell survival upon nutrient-deprived conditions^33,34^, L-2A5HPA increased the survival of cancer cells. Mechanistically, L-2A5HPA enabled glucose flux to be diverted towards the synthesis of aspartate and pyrimidine nucleotides, by maintaining high levels of PC. Aspartate has been described to be an essential metabolite for both bacteria^42,43^ and mammalian cells under hypoxic conditions^44^. Our data showed that aspartate, but not pyruvate, phenocopied L-2A5HPA promoting cell survival under nutrient deprivation conditions. This supports our model in which L-2A5HPA facilitates aspartate biosynthesis under nutrient stress to sustain cell survival.

The de novo synthesis of pyrimidines requires aspartate and functional mitochondria, which are known to undergo degradation via autophagy or mitophagy under nutrient limitation^38^. However, our data demonstrates that the ability of L-2A5HPA to promote cell survival is independent of its capacity to inhibit autophagy. This suggests that the increased availability of aspartate and pyrimidines induced by L-2A5HPA may prevent mitochondrial damage and degradation, increasing the levels of PC and cell survival. Indeed, pyrimidine availability plays a critical role in maintaining mitochondrial integrity. The inhibition of dihydroorotate dehydrogenase (DHODH), a key enzyme in the *de novo* synthesis of pyrimidines, has been shown to trigger mitochondrial damage, and the induction of cell death^45^.

The evidence presented herein also suggests that cancer cells have a stereospecific mechanism to uptake 2A5HPA. Our data reveals that different cancer cells exhibit a variable ability to take up L-2A5HPA, and levels of L-2A5HPA in tumour tissues from different mouse models of liver cancer also differed. Interestingly, the mouse models showing the highest levels of L-2A5HPA are histologically dedifferentiated HCC, which are associated with poorer prognosis^46,47^, highlighting a potential role of L-2A5HPA and malignancy that deserve further investigation.

Although the transporters involved in the uptake of L-2A5HPA are still unknown, and despite the structural similarity between L-2A5HPA and L-serine, no correlation between these metabolites was found in the plasma from patients or *in vitro* studies, which does not support the notion that common transporters for both amino acids exist. Despite this, we observed a strong correlation between the levels of L-2A5HPA, and the levels of pyrimidine biosynthesis intermediates, dihydroorotate and orotate in mouse models of liver cancer, supporting a metabolic link between L-2A5HPA and pyrimidine biosynthesis in an *in vivo* setting.

Interestingly, the patient cohort analyzed in this study received 5-fluorouracil-containing chemotherapy (Folfox and ECX), raising the possibility that L-2A5HPA, by modulating the de novo pyrimidine availability could contribute to the resistance to chemotherapy as has been described by Dong *et al*. ^48^. Further investigation is needed to determine the role of this potential mechanism in the association between L-2A5HPA and patient survival.

The evidence presented here highlights that L-2A5HPA has potential to become a prognostic biomarker for EAC patients treated with pyrimidine analogues-containing chemotherapy, which could prove a valuable tool in combination with other proposed biomarkers such as D-mannose^49^. In addition, we also provide evidence for a biological connection between L-2A5HPA, pyrimidine metabolism and cell survival under nutrient limitations. Although the metabolic adaptation induced by L-2A5HPA partially explains its correlation with overall patient survival, the biological effects downstream of L-2A5HPA seems to be dictated by tumor uptake ability. In summary, the findings presented in this study provide a foundation for future investigations into L-2A5HPA as a prognostic biomarker in esophageal adenocarcinoma (EAC) and underscore its role in modulating tumor metabolism and promoting cell survival.

## Method

### Ethical approvals and clinical cohort information

This study obtained the approval from the School of Medicine ethics committee, University of St Andrews (Ref: MD9202). The samples of plasma EDTA were obtained from the French EsoGastric Tumours (FREGAT) clinical-biological database (NCT 02526095) ^50^ with the approval of the Comité de Protection des Personnes (Ethics Committee) on 10 December 2013, the Agence Nationale de Sécurité du Médicament et des produits de santé (French Health Products Safety Agency) on 13 January 2014, the Comité Consultatif sur le Traitement de l’Information en Matière de Recherche dans le Domaine de la Santé (Advisory Committee on Processing Information Associated with Health Research) on 12 March 2014, and the Commission Nationale Informatique et Libertés (French Data Protection Authority) on 23 December 2014. The samples are preserved in the central BRC of the University hospital of Lille (CRB-CIC1403 du CHU de Lille, BB0033-00030) and the authors are grateful for the work carried out by the entire BRC technical team for this biocollection. All the participants in this study received an information letter and sign a consent form. A cohort of 11 patients who only received pre- and post-operative chemotherapy and have survival data was used for this study. The overall survival was calculated by the difference between the day of plasma sample collection and the event of death of the patients. Although FREGAT clinical-biological database has survival data for 15 patients four of them were discarded as the cause of death was post-operative complication (3) or unknown (1). The correlation analysis between pre-intervention metabolite levels and overall survival has previously been described^51^. The stratification of patients based on their levels of L-2A5HPA was defined as higher or equal and lower compared with the median. Kaplan-Meyer survival plot was used to determine significance between the stratification by L-2A5HPA levels and survival data available from the FREGAT biobank. The description of the patient characteristics is shown in **Extended Data Figure 1**.

### Chemicals and reagents

The chemicals used in this study and their respective suppliers are detailed as follows: 2-amino-5-hydroxypentanoic acid (Biosynth, GAA15289, AstaTech, 51288, BLD Pharmatech Co., Ltd, BL3H9B130671), 2-amino-4methoxybutanoic acid (methoxine, Biosynth, EAA38591), 2-amino-3-hydroxypentanoic acid (Biosynth, CAA28042), 2-amino-4-oxopentanoic acid hydrochloride (Biosynth, XHA23727), NaBH4 (Merck, 452882-5G), L-ethylserinate (ThermoFisher Scientific, AAA1417406), (S)-NIFE (Cayman, Cay17431-5), sodium tetraborate decahydrate (Merck, B9876), sulforhodamine B (Merck, 230162), trichloroacetic acid (Merck, T6399-500G), U-^13^C_6_-D-glucose (Cambridge Isotope Laboratories Inc., CLM-1396-5), aspartic acid (Merck, A9256), sodium pyruvate (Merck, 113-24-6), L-serine (Merck, S4311), rapamycin (Biosynth, AE27685), RIPA buffer (Millipore, 20-188), Laemmli buffer (Bio-Rad, 1610747).

### Synthesis L and D 2A5HPA enantiomers

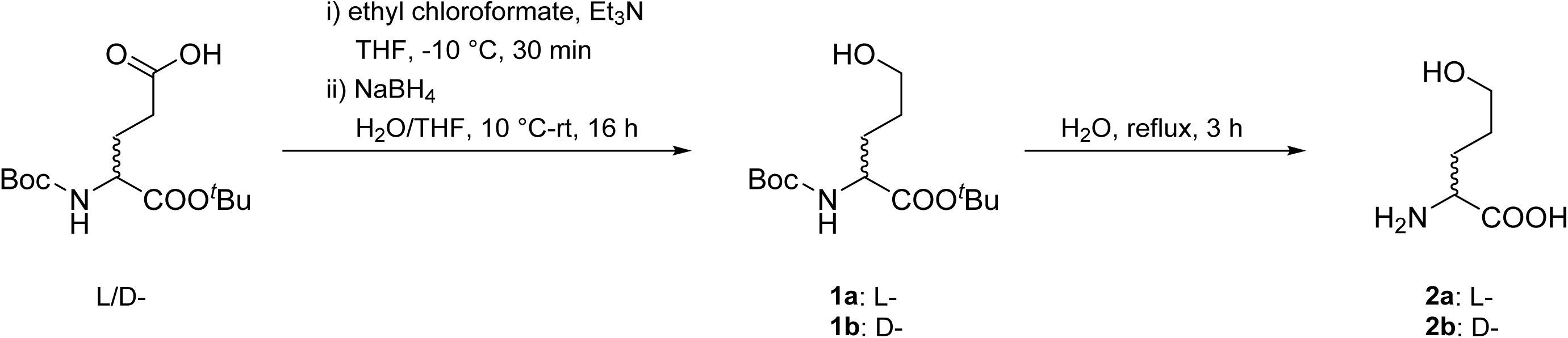

Reduction of Boc-Glu-O*^t^*Bu: An oven-dried flask with a magnetic stir bar was charged with Boc-Glu-O*^t^*Bu (1 eq.). The flask was sealed with a septum and sparged with N_2_. THF (0.25 M, final concentration) and Et_3_N (1.03 eq.) were added using a syringe, and the mixture was cooled to –10 °C. Ethyl chloroformate (1.03 eq.) was added dropwise via syringe, and the resulting mixture was stirred at –10 °C for 30 min. The precipitate was filtered off, and the supernatant was added dropwise to a solution of NaBH_4_ (3 eq.) in a mixture of H_2_O/THF (1:4, 0.2 M final concentration) at 10 °C. The resulting mixture was stirred at room temperature for 16 h. The mixture was acidified with 1 M HCl to pH 3 and extracted with EtOAc (3 × 30 mL). The combined organic extracts were washed with 5% aqueous NaHCO_3_ (2 × 30 mL), saturated aqueous NaCl (2 × 30 mL), dried over Na_2_SO_4_, and concentrated in vacuo. The crude product was purified by column chromatography on silica gel (ethyl acetate/*^n^*hexane, 1:9 → 1:1 v/v).

Cleavage of *^t^*Bu and Boc groups: A mixture of Boc-Nva(5-OH)-O*^t^*Bu 1a-b (1 eq.) in H_2_O (0.05 M) was heated under reflux for 3 h. The mixture was allowed to cool to room temperature and then concentrated in vacuo to obtain the product as a white solid. Boc-L-Nva(5-OH)-O*^t^*Bu (**1a**) : Prepared following General procedure 1 using Boc-L-Glu-O*^t^*Bu (2.002 g, 6.60 mmol, 1 eq.), ethyl chloroformate (0.65 mL, 6.8 mmol, 1.03 eq.), and Et_3_N (0.95 mL, 6.8 mmol, 1.03 eq.) in THF (25 mL); and then NaBH_4_ (0.749 g, 19.80 mmol, 3 eq.) in a mixture of H_2_O (1.7 mL) and THF (6.8 mL). Purification by silica gel column chromatography (ethyl acetate/n-hexane, 1:9 → 1:1 v/v) afforded product **1a** as a colorless oil in 73% yield (1.389 g, 4.80 mmol). ^1^H NMR (500 MHz, DMSO) δ 7.07 (d, *J* = 7.7 Hz, 0.8H), 6.73 (d, *J* = 7.7 Hz, 0.2H), 4.41 (t, *J* = 5.1 Hz, 1H), 3.75 (td, *J* = 8.5, 5.4 Hz, 0.8H), 3.67 (m, 0.2H) 3.36 (m, 2H), 1.73 – 1.58 (m, 1H), 1.58 – 1.47 (m, 1H), 1.46 – 1.31 (m, 18H); ^13^C NMR (126 MHz, DMSO) δ 171.9, 155.6, 80.1, 78.0, 60.2, 54.3, 28.9, 28.2, 27.7, 27.4. This data is consistent with literature values ^52^.

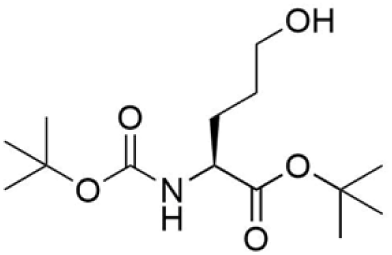

Boc-D-Nva(5-OH)-O*^t^*Bu (**1b**): Prepared following general procedure 1 using Boc-D-Glu-O*^t^*Bu (1.517 g, 5.00 mmol, 1 eq.), ethyl chloroformate (0.49 mL, 5.15 mmol, 1.03 eq.), and Et_3_N (0.72 mL, 5.15 mmol, 1.03 eq.) in THF (19 mL); and then NaBH_4_ (0.567 g, 15.00 mmol, 3 eq.) in a mixture of H_2_O (1.3 mL) and THF (5.0 mL). Purification by silica gel column chromatography (ethyl acetate/*^n^*hexane, 1:9 → 1:1 v/v) afforded product **1b** as a colorless oil in 66% yield (0.956 g, 3.30 mmol). ^1^H NMR (500 MHz, DMSO) δ 7.07 (d, *J* = 7.7 Hz, 0.8H), 6.73 (d, *J* = 7.7 Hz, 0.2H), 4.41 (t, *J* = 5.1 Hz, 1H), 3.75 (td, *J* = 9.1, 5.3 Hz, 0.8H), 3.67 (m, 0.2H), 3.36 (m, 2H), 1.72 – 1.59 (m, 1H), 1.58 – 1.48 (m, 1H), 1.44 – 1.31 (m, 18H); ^13^C NMR (126 MHz, DMSO) δ 171.9, 155.5, 80.1, 78.0, 60.2, 54.3, 28.9, 28.2, 27.7, 27.5. This data is consistent with literature values ^52^.

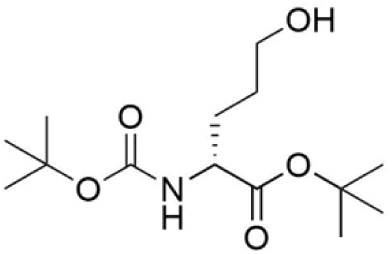

H-L-Nva(5-OH)-OH (**2a**): Prepared following general procedure 2, starting from **1a** (0.351 g, 1.21 mmol). Product **2a** isolated as a white solid in 97% yield (0.157 g, 1.18 mmol). ^1^H NMR (500 MHz, D_2_O) δ 3.77 (t, J = 6.13 Hz, 1H), 3.63 (m, 2H), 2.01 – 1.84 (m, 2H), 1.73 – 1.55 (m, 2H); ^13^C NMR (126 MHz, D_2_O) δ 174.6, 60.9, 54.5, 27.2, 27.1. This data is consistent with literature values ^21–23^.

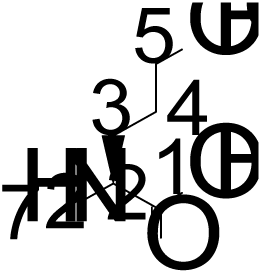

H-D-Nva(5-OH)-OH (**2b**): Prepared following General procedure 2, starting from **1b** (0.492 g, 1.70 mmol). Product **2b** was isolated as a white solid in 98% yield (0.222 g, 1.67 mmol). ^1^H NMR (500 MHz, D_2_O) δ 3.76 (t, *J* = 6.12 Hz, 1H, H_2_), 3.63 (m, 2H, H_5_), 1.99 – 1.83 (m, 2H, H_3_), 1.63 (m, 2H, H_4_); ^13^C NMR (126 MHz, D_2_O) δ 174.6, 60.9, 54.5, 27.2, 27.1. This data is consistent with literature values ^21–23^.

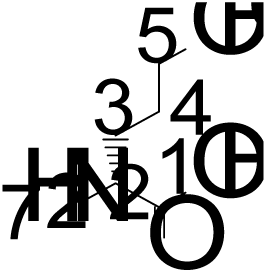

### Synthesis of 2-amino-4-hydroxypentanoic acid

For the synthesis of 2-amino-4-hydroxypentanoic acid, 2 mg of 2-amino-4-oxopentanoic acid hydrochloride (Biosynth, XHA23727) were dissolved in 200 µL of methanol, and 2 mg of NaBH_4_ (Merck, 452882-5G) were added to the solution and incubated for 2 h on ice ^53^. Finally, 800 µL of water was added to the solution. The final product was diluted in extraction buffer to obtain a final concentration of 5 µM and subsequent analysis by LC-MS and LC-MS/MS.

### Cell Culture

U2OS and U2OS GFP-LC3 cells were kindly provided by Dr. Saverio Tardito (Cancer Research UK Scotland Institute, Glasgow UK). Esophageal cancer cells OAC-P4C and SK-GT-4 were kindly provided by Prof. Neil Carragher (Institute of Genetics and Cancer, the University of Edinburgh, UK). The SW982 cells were kindly provided by Dr. Javier Martin-Broto (University Hospital Fundacion Jimenez Diaz, Spain). The cell lines were cultured in DMEM (Corning, 10-013-CV) supplemented with 10% FBS (U2OS, U2OS GFP-LC3, SW982, OAC-P4C and SK-GT-4).

Metabolic analysis of the esophageal cancer cells (OAC-P4C and SK-GT-4) was performed by culturing the cell lines in MEM (Gibco, 21090-22) supplemented with 10% FBS, 2 mM glutamine (Gibco, 25039-081), 1× non-essential amino acids (Gibco, 11140-035) and 1 mM sodium pyruvate (Merk 58636), and at 37 °C, 5% CO2 in a humidified atmosphere. The OAC-P4C and SK-GT-4 cells were seeded as 5× 10^5^ cells per well on 6-well plates and after 24 hours, the cells were incubated in Plasmax^TM^ (Ximbio, 156371) supplemented with 2.5% FBS for additional 24 hours at 37 °C, 5% CO_2_ in humidified atmosphere. All the cells used in the manuscript were mycoplasmas tested every 5-months using a PCR kit (#G238, ABM or Venor^(R)^ GeM onStep, Minerva biolabs).

### Nutrient deprivation conditions

The autophagy and nutrient deprivation experiments were performed in EBSS (Gibco, 24010043). U2OS or SW982 were seeded at the density of 2×10^5^ in a 6 well plate. 24 h after, the cells were then exposed to nutrient deprivation conditions (EBSS) in the presence or absence of 0.5 to 2 mM L-2A5HPA, 2 mM D-2A5HPA, 2 mM L-serine, 2 mM L-aspartate or 2 mM pyruvate for 4 to 5 days at 37 °C, 5% CO2 in a humidified atmosphere.

### Recovery experiments

To assess the ability of cells to recover after 4 to 5 days on starvation, cells were incubated in complete DMEM in the absence of any compounds for a further 5 to 8 days. Then the cells were fixed and stained with sulforhodamine B as described in Ackermann et al. (2024). Briefly, after their respective treatment, cells were washed twice with PBS, fixed with 3% TCA containing 1% acetic acid for 10 min. Thereafter, cells were stained with 0.057% sulforhodamine B (SRB) in 1% acetic acid in water for 10 min. The excess of sulforhodamine B was removed by washing the cells with 1% acetic acid in water twice, and plates were air dried. The quantification of SRB was performed redissolving it on 10mM Tris-HCl pH=8 and measuring the absorbance at 510 nm using the CLARIOstar Plus Microplate Reader.

### Western blotting

Cells were lysed in RIPA buffer containing mini-Complete protease inhibitor cocktail and phosphatase inhibitors (ThermoFisher Scientific, A32961). Total protein concentration was quantified using a Pierce BCA kit (Thermo Fisher Scientific, 23227). Equal amounts of protein were heated at 95 °C for 10 min and separated (100 V) in 9,5%, 11% or 16% acrylamide/bisacrylamide gels for SDS-PAGE. Proteins were transferred onto nitrocellulose membranes (Thermo Fisher Scientific, 21882), then blocked for one hour in 5% skimmed milk in TBS. Thereafter, the membranes were incubated overnight at 4 °C with the primary antibodies. The primary antibodies used: Cleaved Caspase 3 (Cell Signaling, 9662 dilution 1:500), Cleaved PARP (Cell Signaling, 9532 dilution 1:1000), P62 (Cell Signaling, 397495 dilution 1:1000), LC3BII (Cell Signaling, 3868 dilution 1:1000), P-S6 S235/236 (Cell Signaling 4858 dilution 1:1000), S6 (Cell Signaling, 2317 dilution 1:1000), β-actin (Cell Signaling 4970S dilution 1:1000), Vinculin (Merck, MAB3574 dilution 1:1000), PCB (ab126707 dilution 1:1000). Secondary antibodies HRP-linked antibodies: Anti-rabbit IgG HRP-linked Ab (Cell Signaling Cat#7074, dilution 1:1000) and Anti-mouse IgG HRP-linked Ab (Cell Signaling Cat#7076, dilution 1:1000). Secondary antibodies were incubated for 1 h at room temperature. The blots were revealed with Clarity Western ECL Substrate (Bio-Rad, 1705061) and imaged with the iBright F1500 (Thermofisher). All uncropped and unprocessed images are presented in **Extended Data Figure 3**.

### LC-MS metabolomics

Metabolites from the plasma, cells and tissue were extracted using the metabolomic extraction buffer (5:3:2 ratio of methanol, acetonitrile, water) ^55^. 5 µL of plasma was transferred into 250 µL of extraction buffer in an Eppendorf tube. The samples were centrifuged at 18,000 rpm for 10 min at 4° C, and the supernatant was collected into mass spectrometry vials and stored at -80 ° C. All cell lines used were seeded at 5×10^5^ cells/well in 6-well plates, and after the respective treatment, 500 µL of extraction buffer was added into each of the wells and incubated at 4 °C for 5 min. The supernatant was collected into a fresh Eppendorf tube and centrifuged at 18,000 rpm for 10 min at 4 ° C. Finally, the supernatants were collected into mass spectrometry vials. Protein concentration in each of the wells was quantified using a modified Lowry method ^55^.The samples were normalized to the protein concentration in each of the wells. Finally, 20 mg of tissue was disrupted in 1 ml of extraction buffer using the Precellys 24 Tissue Homogenizer (Bertin Technologies). The extracts were centrifuged at 18,000 rpm for 10 min to remove proteins and cell debris. The supernatant was collected into mass spectrometry vials and stored at -80 °C for subsequent analysis^55^. The tumor tissue was normalized by weight, maintaining the same weight/extraction buffer ratio.

The extracts were injected (5µL) and separated on a ZIC-pHILIC column (SeQuant; 150 mm × 2.1 mm, 5 μm; Merck) coupled to a ZIC-pHILIC guard column (SeQuant; 20 mm × 2.1 mm) using an Ultimate 3000 HPLC system (ThermoFisher Scientific).

A 15-min linear gradient was used for the chromatographic separation at constant temperature (45 °C), starting with 20% ammonium carbonate (20 mM, pH 9.2) and 80% acetonitrile, decreasing acetonitrile until 20% with a constant flow rate of 200 μl min^−1^. The Q Exactive Orbitrap mass spectrometer (ThermoFisher Scientific) equipped with electrospray ionisation was coupled to the HPLC system. The polarity switching mode was used for metabolite profiling, a resolution (RES) of 35,000 or 70,000 at 200 m/z to enable both positive and negative ions to be detected across a mass range of 75 to 1,000 m/z (automatic gain control (AGC) target of 1 × 10^6^ and maximal injection time (IT) of 250 ms).

Data-independent fragmentation was performed to acquire fragmentation spectra of specific metabolites including 2A5HPA (positive polarity, m/z 134.0811). Fragmentation spectra were continuously recorded with the following parameters: 17,500 RES, isolation width of 0.7 m/z, AGC target of 1 × 10^5^, max IT of 250 ms and stepped normalized collision energy of 25, 60 and 95.

### Cellular uptake of L-2A5HPA and D-2A5HPA

To examine the cellular uptake of L-2A5HPA uptake U2OS, SK-GT-4 and OA-P4C were seeded at 3×10^5^ in 6-well plates and incubated for 24h with complete Plasmax^TM^ and then treated with L-2A5HPA 50 µM for 1 min. The uptake of D-2A5HPA was assessed in similar conditions in U2OS cells. The metabolites were extracted as described in the LC-MS metabolomic section.

### (S)-NIFE derivatization

The (S)-NIFE derivatization reaction was prepared using a modified protocol from Vessel et al ( 2011). For this, 100 µL of pooled plasma from patients was combined with 500 µL of acetonitrile in an Eppendorf tube and vortexed for 5 s. To remove protein precipitates, the tube was centrifuged at 13,000 rpm for 5 min, and the supernatant was collected into a fresh Eppendorf tube. The supernatant was evaporated using a Vacuum concentrator (Savant, SC210A Speed Vac concentrator), and resultant metabolites were resuspended in 50 µL of milliQ water. The chemically pure synthesized enantiomers L- and D-2A5HPA were dissolved in water to a final concentration of 50 µM. The derivatization reaction was performed using either 50 µL of resuspended plasma, 50 µL L-5H2APA (50 µM) or 50 µL D-5H2APA (50 µM) and 25 µL of tetraborate 0.2 M in water and 25 µL of (S)-NIFE (2mg/mL in acetonitrile). This mixture was shaken by an orbital shaker (500 rpm) for 20 min and then analyzed by LC-MS.

### LC-MS of (S)-NIFE-L-2A5HPA and (S)-NIFE-D-2A5HPA

Analysis of (S)-NIFE derivatized D- or L-2A5HPA was performed using a Q-Exactive mass spectrometer (ThermoFisher Scientific) coupled to an Ultimate 3000 HPLC system. Chromatographic separation was achieved using a ZORBAX Eclipse plus C18 column (2.1×50mm 1.8micron, PN 959757-902) using water and acetonitrile with 0.1% formic acid as mobile phase A and B, respectively. Gradient elution for the diastereomers was as follows: 0 min, 99% A; 23 min 85% A; 25 min 10% A; 27 min 10% A; 27 min 99% A; 30 min 99% A. Column temperature was kept at 37 °C, with a flow rate of 0.350 mL/min. Metabolite profiling was conducted in positive electrospray ionization mode, using a resolution of 70,000 at 200 m/z. The mass spectrometer was set to scan a mass range of 100 to 1,000 m/z, with an automatic gain control (AGC) target of 1 × 10⁶ and a maximum injection time (IT) of 250 ms.

### U-^13^C_6_-glucose tracing

To investigate changes in glucose metabolism flux, we performed a U-^13^C_6_-glucose tracing experiment. U2OS cells were seeded at a density of 3×10^5^ in 6-well plates in complete media. After 24 h, the culture media was replaced with glucose-free EBSS supplemented with 1g /L of U-^13^C_6_-glucose. The cells were incubated with this media in the absence or presence of 2 mM L-2A5HPA for 72 h. The metabolites were extracted as described in the LC-MS metabolomic section.

### Metabolomics data analysis

The targeted metabolomic analysis was carried out using an in-house library of 80 metabolites using their m/z and retention time. The quantification of the peak area was performed using Skyline 24.1.0.199. The untargeted metabolomics analysis was performed using Compound Discoverer v.3.1, and the identification was based on both a publicly available database (MS/MS data) and an in-house standards library with specific mass and retention time. In addition, we used Sirius v6.1.0 to support the structure elucidation ^56^.

### Fluorescence microscopy

To assess autophagy we used U2OS GFP-LC3 cells^33,34^. These cells were seeded at 2×10^5^ cells/well in a 12 well plate, grown on coverslips and starved with EBSS in the presence or absence of L-2A5HPA for 5 days. Thereafter, cells were fixed with 4% paraformaldehyde in PBS for 15 min at room temperature. U2OS GFP-LC3 cells were then mounted after the fixation with Prolong containing DAPI (P36966, Invitrogen). The slides were imaged using a Leica DM55000 B fluorescence microscope (Leica).

### Animal samples

All animal experiments were performed in accordance with UK Home Office licenses (PP0604995) and in accordance with the UK Animal (Scientific Procedures) Act 1986 and EU Directive 2010. The protocols were also reviewed by the Animal Welfare and Ethical Review board of the University of Glasgow. Welfare of animals was defined by clinical symptoms, including visible masses, any degree of reduced mobility/distress, weight loss or evidence of hemorrhage. However, no maximal tumor volume end points for intrahepatic tumors were mandated. No mouse exceeded the humane endpoints stipulated in the Home Office Licenses. All the tumor and tissue samples were provided by Prof. Tom Bird as previously described ^40^.

All the samples were collected from male mice on a mixed background. The following transgenic mice strains were used: Ctnnb1tm1Mmt^(Ctnnb1ex3)^, Gt(Rosa)26Sor^tm1(MYC)Djmy^ (R26LSL-MYC), Trp53^tm1Brn^ (Trp53fl), Trp53^tm2Tyj^ (Trp53R172H), Cdkn2a^tm1.1Brn^ (Cdkn2a^KO^), Pten^tm2Mak^ (Pten^fl^), Axin1^fl40^. Genotyping was performed by Transnetyx. Mice were induced between 8 and 12 weeks of age with AAV8.TBG.PI.cre.rBG (AAV8-TBG-cre) (Addgene, 107787-AAV8). Heathy aged-matched controls were induced with a mock virus (AAV8.TBG.PI.Null.bGH or AAV8-TBG-Null) (Addgene, 105536-AAV8). The mice received a dose of 2 × 10^11^ (TP53/Myc model), 6.4×10^8^ GC or 1.28×10^8^ GC per mouse in PBS as previously described ^40^.

In this study 3-6 mice per genotype were sampled at the end point, which is defined as the mouse having reached a liver weight/body weight ratio of 20% or exhibiting side effects from the tumour, such as tumour haemorrhaging. Macroscopic tumours were snap-frozen on dry ice and stored at -80°C.

### Statistical analysis

The statistical analysis of the metabolomics data, correlations and other comparisons were carried out using GraphPad Prism v10 (t-test, one-way ANOVA, Pearson correlations and ROC).

Software: Compound Discoverer v.3.1 was used for the untargeted metabolomic analysis. Skyline 25.1.0.237 was used for the targeted metabolomics analysis. Xcalibur version 4.3 was used to obtain the extracted ion chromatograms and the ms^2^ spectra. Sirius 6.1.0 was used to support structure elucidation of the unknown metabolite.

## Authors Contributions

All authors have read and approved the final version of this manuscript. Author contributions: Conceptualization: M.E.K., V.H.V. Visualization M.E.K., V.H.V. Resources H.A., K.M.R., R.D., D.S.M., S.A., S.M., M.M., T.M.D., A.G., T.G.B., M.H., G.P., A.J.S., A.J.B.W., D.S. Methodology: A.H.U., M.C., A.J.B.W., D.S. Formal analysis: M.E.K., A.H.U., M.C., K.B.H., V.H.V. Performing experiments: M.E.K., K.B.H., M.C., H.A. Data Curation: M.F.F. Writing - Original Draft: M.E.K., V.H.V. Writing - Review & Editing: K.M.R., R.D., D.S.M., T.G.B., M.H., G.P., A.J.S., A.J.B.W., D.S., V.H.V. Supervision: T.G.B., M.H., G.P., A.J.S., A.J.B.W., D.S., V.H.V. Funding acquisition: S.A., T.G.B., A.J.S., A.J.B.W., D.S., V.H.V.

## Acknowledgments

We thank the FREGAT Working Group, the Data Treatment Centre for administering the FREGAT database and providing access to the data, as well as the Biological Resource Centres and Tumor Banks participating in the FREGAT project for their valuable contributions.

---

**Extended Data Figure 1.**
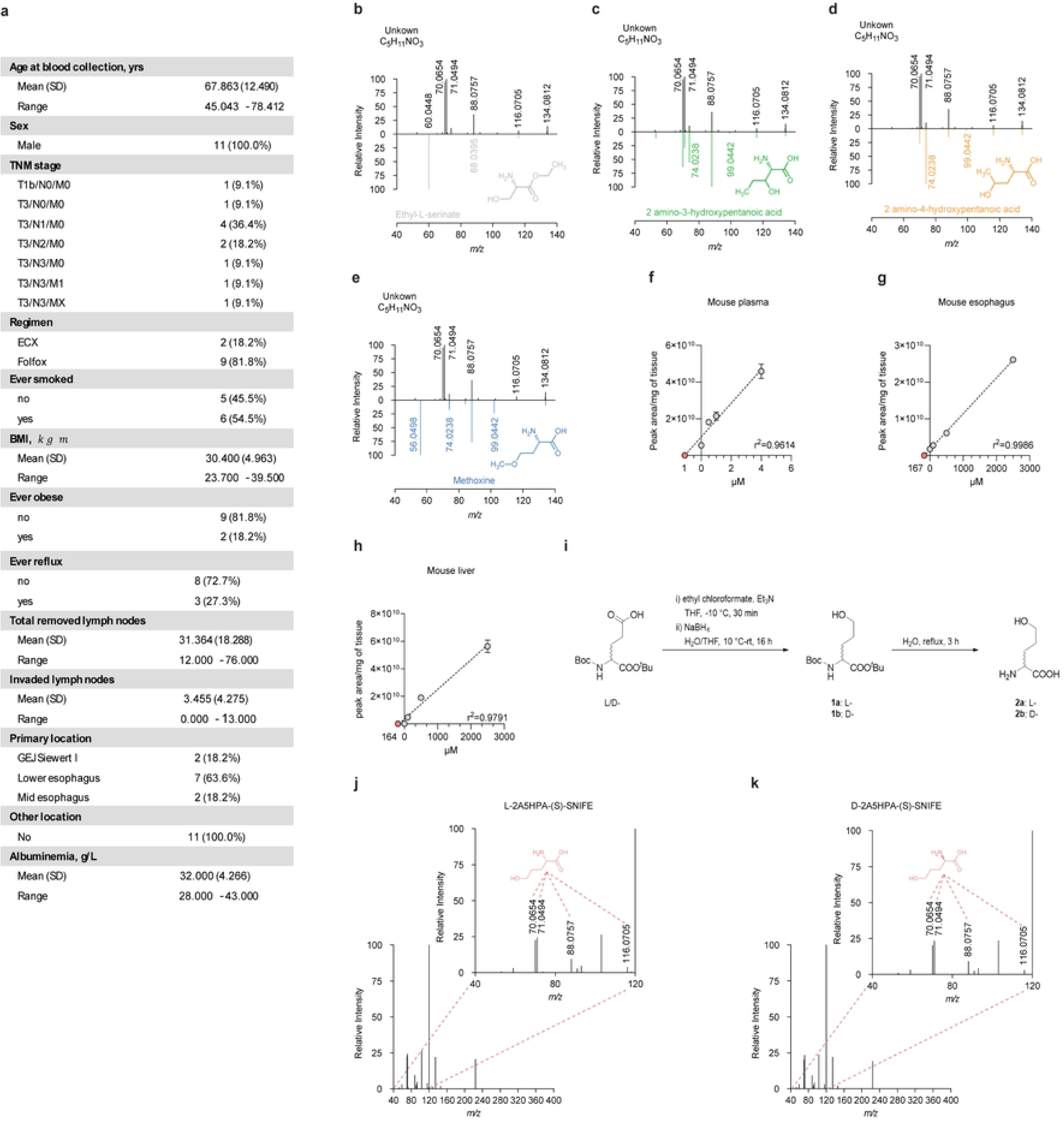
Fragmentation pattern comparison between the unknown C_5_H_11_NO_3_ and the potential chemical structures, and the estimation of L-2A5HPA physiological concentration in mouse plasma and tissues. **a**) Characteristics of the eleven patients selected for this study. Mirror plots showing the fragmentation pattern between the unknown C_5_H_11_NO_3_ from pooled plasma samples (black) and ethyl-serinate (grey, **b**), unknown C_5_H_11_NO_3_ and L-2-amino-3-hydroxy-pentanoic acid (green, **c**), unknown C_5_H_11_NO_3_ and 2-amino-4-hydroxyl-pentanoic acid (orange, **d**), unknown C_5_H_11_NO_3_ and methoxine (blue, **e**). Absolute quantification of L-2A5HPA using the standard addition method in pooled mouse plasma (n=3) (**f**), mouse esophagus tissue (n=1) (**g**), and mouse liver tissue (n=1) (**h**). **i**) Schematic for the organic synthesis of L-2A5HPA and D-2A5HPA. Fragmentation analysis of the (S)-NIFE-derivatised L-2A5HPA (**j**) and (S)-NIFE-derivatised D-2A5HPA (**k**). Insets show the fragments of L and D-2A5HPA. Circles represent the average of technical duplicates.

**Extended Data Figure 2.**
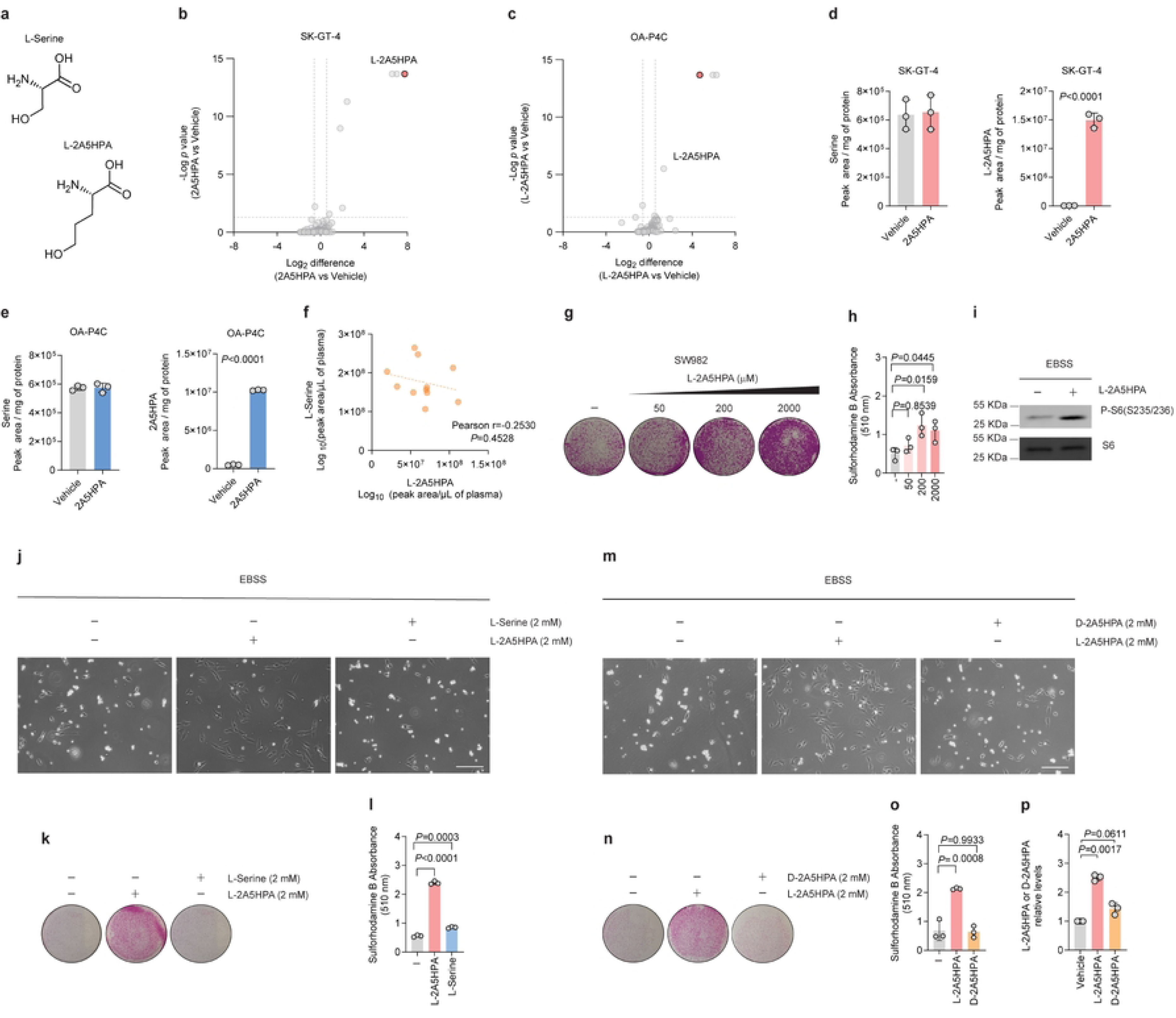
Untargeted metabolomic analysis of esophageal cancer cells, and L-2A5HPA effect on survival and mTORC1 activity. **a**) Chemical structure of L-serine and L-2A5HPA. Representative volcano plot of untargeted analysis comparing SK-GT-4 (**b**) and OA-P4C cells (**c**) treated with or without L-2A5HPA (50 µM). The grey circles represent metabolites detected, and L-2A5HPA is highlighted in red. Intracellular levels of L-serine and L-2A5HPA are shown in SK-GT-4 cells (**d**) and OA-P4C cells (**e**). **f**) Correlation between the plasma levels of serine and L-2A5HPA (n=11). The Pearson r value and *P* value are shown. **g**) Representative image of SW982 cells that were nutrient-deprived in the absence or presence of L-2A5HPA (0, 50, 200 and 2,000 µM) for 4 days, recovered in complete media for 8 days and stained with sulforhodamine B. **h**) Quantification of sulforhodamine B absorbance at 510 nm. **i**) Immunoblot analysis of U2OS cells nutrient-deprived in the presence or absence of L-2A5HPA for 5 days. mTORC1 activation was measured by the levels of phosphor-S6 (S235-236). Total S6 is shown as a loading control. **j**) Representative images from U2OS cells starved and treated with vehicle, L-Serine (2mM) or L-2A5HPA (2mM) for 5 days. **k**) Representative images from cells treated as in **j**), then exposed to complete media for 5 days and stained with sulforhodamine B. **l**) Quantification of sulforhodamine B absorbance at 510 nm from three biological replicates. **m**) Representative images of starved cells treated with either L-2A5HPA (2mM) or D-2A5HPA (2mM) for 5 days. **n**) Representative images from cells stained with sulforhodamine B after being treated as in **m**), then exposed to complete media for 5 days. **o**) Quantification of sulforhodamine B absorbance at 510 nm of three biological replicates. **p**) Relative intracellular levels of L-2A5HPA or D-2A5HPA in U2OS cells after being exposed to 50 µM of each of the enantiomers for 1 min. Bars represent the average ± SEM of three independent replicates. Data were analyzed using a one-way ANOVA, Student’s t-test or One-sample t-test (p). Circles represent metabolites, independent experiments or individual patients.

**Extended Data Figure 3.**
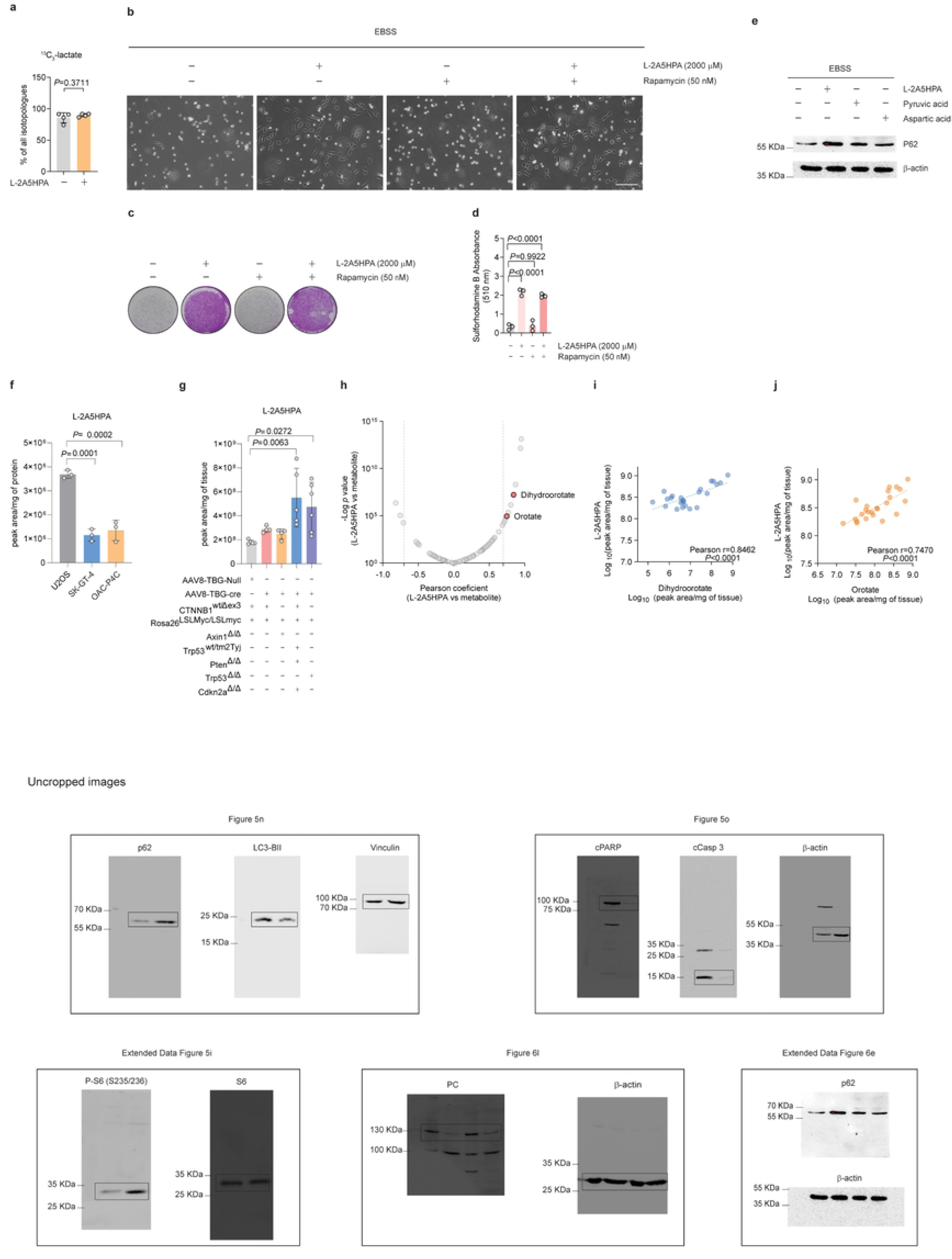
^13^C_6_-Glucose tracing, L-2A5HPA uptake in cell lines and the levels of L-2A5HPA in tumour tissue from different mouse models of liver cancer. **a)** Enrichment of ^13^C_3_-lactate from ^13^C_6_-glucose in U2OS cells starved in the presence or absence of L-2A5HPA for 72h. **b**) Representative image of U2OS cells starved for 5 days with or without L-2A5HPA in the presence or absence of rapamycin (50nM). **c**) Representative image of U2OS cells treated as in **b**) and then exposed to complete media for 5 days and stained with sulforhodamine B. **d**) Quantification of sulforhodamine B absorbance at 510 nm. **e**) Immunoblot analysis of p62 in U2OS cells starved for 5 days in the absence of any compound and in the presence of either L-2A5HPA, L-aspartate or pyruvate. β-actin was used as a loading control. **f**) Intracellular levels of L-2A5HPA in U2OS, SK-GT-4 and OAC-P4C cells after being exposed to 50 µM L-2A5HPA for 1 min. **g)** Relative levels of L-2A5HPA in tumour tissue from different genetically engineered mouse models of hepatocellular carcinoma ^40^. **h**) Correlation between levels of L-2A5HPA and all the other metabolites detected in tumour and healthy tissue. Dihydroorotate and orotate are highlighted in red. Correlation between L-2A5HPA and dihydroorotate levels (n=24) (**g**) or orotate levels (n=24) **(h**) in tumour and healthy tissue. The Pearson coefficient and P values are shown in the graphs. **i**) Uncropped blot images corresponding to the western blots presented in the main and supplementary figures are provided. Each image includes the figure in which they are presented and the antibodies used. Bars represent the average ± SEM. Data were analyzed using a one-way ANOVA or Student’s t-test. Circles represent independent experiments, metabolites or tumour and healthy tissue sampled from individual animals.

